# Attentional engagement with target and distractor streams predicts speech comprehension in multitalker environments

**DOI:** 10.1101/2025.04.04.647157

**Authors:** Alice Vivien Barchet, Andrea Bruera, Jasmin Wend, Johanna M. Rimmele, Jonas Obleser, Gesa Hartwigsen

## Abstract

Understanding speech while ignoring competing speech streams in the surrounding environment is challenging. Previous studies have demonstrated that attention shapes the neural representation of speech features. Attended streams are typically represented more strongly than unattended ones, suggesting either enhancement of the attended or suppression of the unattended stream. However, it is unclear how these complementary processes support attentional filtering and speech comprehension on different hierarchical levels. In this study, we used multivariate temporal response functions to analyze the EEG signals of 43 young adults (24 women), examining the relationship between the neural tracking of acoustic and higher-level linguistic features and a fine-grained speech comprehension measure. We show that the neural tracking of word and phoneme onsets and word-level linguistic features in the attended stream predicted comprehension at the individual single-trial level. Moreover, acoustic tracking of the ignored speech stream was positively correlated with comprehension performance, whereas word level linguistic neural tracking of the ignored stream was negatively correlated with comprehension. Collectively, our results suggest that attentional filtering during speech comprehension requires target enhancement as well as distractor suppression at different hierarchical levels.

**Significance Statement:** In social settings, speech comprehension is often challenged by the presence of multiple speakers talking simultaneously. The ability to focus on a relevant stream while ignoring irrelevant speech information in the background is crucial for successful and efficient interpersonal interactions. However, the precise neural mechanisms underlying this selective filtering process remain unclear. We establish the interplay of acoustic and higher-level information as objective markers of attentional selection and comprehension success.

## Introduction

In everyday life, spoken speech streams are often obscured by background noise or competing speech streams, resulting in complex listening conditions. To successfully manage these challenges, listeners rely on information from multiple hierarchical levels to filter the relevant signals and facilitate comprehension. In this EEG study, we examined how the processing of acoustic and higher-level linguistic features of both attended and unattended speech streams contributes to speech comprehension in a multitalker environment.

Speech is hierarchically organized into multiple levels of information (Berwick et al., 2013; Ding et al., 2016). Neurally, this organization is implemented by low-frequency cortical activity that aligns with the speech signal (Henry & Obleser, 2012; Obleser & Kayser, 2019; Schroeder et al., 2010). This phenomenon, termed neural tracking, has been demonstrated across acoustic, phonemic, and semantic levels (Brodbeck et al., 2018; Broderick et al., 2018; Donhauser & Baillet, 2020; Gillis et al., 2021; Heilbron et al., 2022). Comprehension in challenging listening environments relies on selective attention.

Selective attention is required when multiple stimuli compete for cognitive processing and processing and attentional filtering is needed to select the relevant stream (Moore & Zirnsak, 2017). This selection can be implemented using two mechanisms: target enhancement and distractor suppression (Orf et al., 2023). Attention has been shown to selectively enhance the neural tracking of acoustic properties of the attended stream (Ding & Simon, 2012; Golumbic et al., 2013; Horton et al., 2014; Mesgarani & Chang, 2012; O’sullivan et al., 2015). However, the presence of acoustic neural tracking alone does not indicate comprehension as it also occurs for non-speech sounds and unintelligible speech (Hämäläinen et al., 2012; Karunathilake et al., 2023; Steinschneider et al., 2013; Tierney & Kraus, 2015). Instead, the neural tracking of higher-level linguistic information has been proposed as a neural marker of speech comprehension (Gillis et al., 2021; Verschueren et al., 2022).

This is supported by evidence showing that higher-level features are typically not neurally represented for unattended speech (Brodbeck et al., 2018; Broderick et al., 2018). Previous research has indicated that attentional selection in competing speech situations involves higher-order, top-down mechanisms (Golumbic et al., 2013; Rimmele et al., 2015). Thus, higher level distractor suppression could be an important mechanism supporting attentional selection. Neural tracking of acoustic information has been observed for ignored speech streams (Ding & Simon, 2012; Golumbic et al., 2013; Horton et al., 2014; Kaufman & Golumbic, 2023; O’sullivan et al., 2015; Rimmele et al., 2015). However, its relation to comprehension remains unclear, as the tracking of distractor acoustic information has been shown to be unrelated (Orf et al., 2023) or even positively related to speech comprehension performance (Fiedler et al., 2019).

Direct evidence for the interplay of lower- and higher-level neural representations of attended and unattended speech streams in predicting speech comprehension is lacking. Many previous studies using continuous listening paradigms were unable to capture rapid variations in comprehension success due to the lack of detailed behavioral responses (Brodbeck et al., 2018; Broderick et al., 2018; Gillis et al., 2021; Verschueren et al., 2022). In this study, we employed a fine-grained, temporally resolved measure of comprehension (i.e., sentence repetition) to provide deeper insights into the dynamics of speech comprehension. Additionally, we quantified acoustic and linguistic speech representations for attended and unattended speech streams in a fine-grained fashion.

Given the previous evidence, we expected that target enhancement, as indexed by enhanced processing of the attended speech stream at acoustic and linguistic levels, would be positively correlated with comprehension performance. For the unattended stream, we expected that distractor suppression at the acoustic level would not be related to comprehension (see Orf et al., 2023). In contrast, we expected that distraction by the unattended stream would manifest in an increased higher-level tracking of the unattended stream. Therefore, linguistic tracking of the distractor stream was expected to be negatively related to comprehension performance.

## Materials and Methods

### Participants

The sample included N = 43 right handed German native speakers aged 19 to 51 (M = 33.86, SD = 10.57; 24 women). Participants had normal hearing and no history of neurological or psychiatric disorders. Normal hearing abilities were confirmed by a pure tone audiometry prior to the experiment. Normal hearing was defined as having a mean pure tone audiometric threshold of below 25 dB across a frequency range of 250 to 8000 Hz in at least one ear. The mean pure tone threshold for each participants’ better ear in our sample was M = 4.31 dB (range = 0 - 15.5 dB; see Figure S1 for the full audiometric results). The average difference between the better ear and the other ear was 3.3 dB (range = 0 - 14 dB).

Participants were recruited from the participant database at the Max Planck Institute for Human Cognitive and Brain Sciences and received monetary compensation of 12 *€* per hour for participation. The study was approved by the ethics committee of the Medical Faculty at Leipzig University (ethics vote: 032/24-ek).

### Procedure

Participants listened to 240 trials separated into 12 blocks of 20 sentences each. Each trial consisted of two sentences presented in parallel. One sentence was spoken by a female speaker and the other one was spoken by a male speaker. Participants were instructed to follow the male speaker while ignoring the female speaker. Thus, all repeated target sentences were spoken by the same male speaker. After listening to the sentences and a brief period of 1 second to avoid EEG artifacts, participants had to repeat the target sentence. If participants did not understand the complete sentence, they were instructed to repeat the portion of the sentence they did understand. If they were unable to understand any part of the sentence, they were asked to say so. The behavioral paradigm is illustrated in figure 1. Experiment control was administered by the Psychophysics Toolbox Version 3 running on Matlab on a Windows machine (Brainard & Vision, 1997; Kleiner et al., 2007). The experiment was conducted in a sound-proof, electromagnetically shielded booth. Stimuli were presented using Koss KSC75 headphones clipped to the participants’ ears. Speech was recorded using a high quality condenser microphone (Rode NT55) at 44100 Hz. Target sentences were presented at six individually adjusted target-to-distractor ratios (T/D) [-2, -1, 0, 1, 2, 10 dB relative to their 50% speech reception threshold]. We additionally calculated word-by-word audibility by quantifying the proportion of the glimpsed target signal (see below). The latter condition served as a control condition, in which the target sentence should be well comprehended by every participant and thus serving as a control for attention and working memory. Control trials were excluded from behavioral and neural analyses and mean accuracy in the control condition was added as a control variable in the regression analysis (see below). Accuracy in the control condition was high (M = 97%, range = [93 - 99 %]), indicating that all participants attentively completed the task.

**Figure 1:**
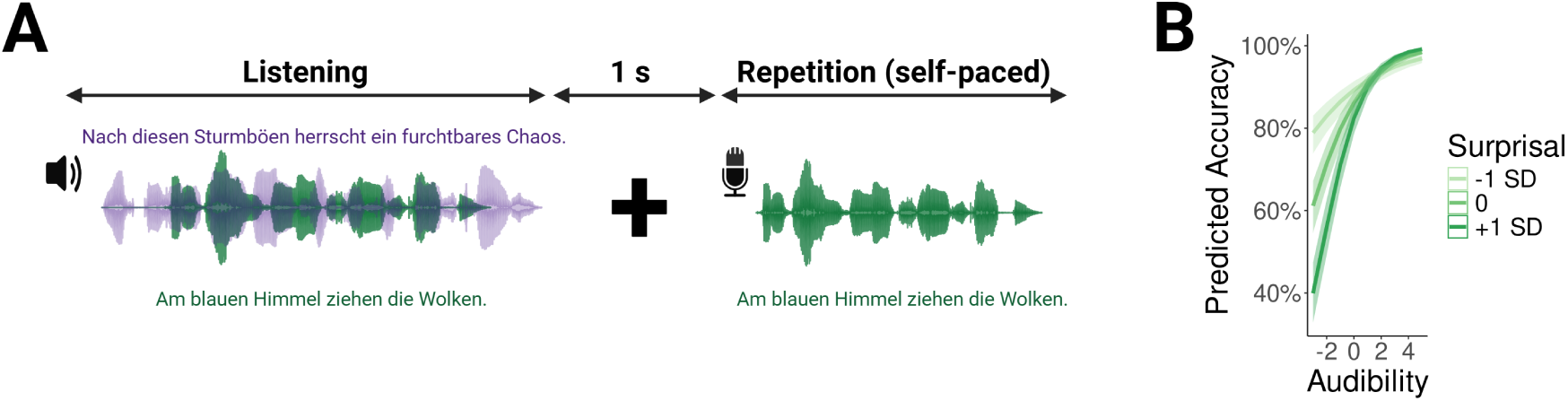
Behavioral analysis. A: Behavioral Paradigm. Participants listened to 240 trials comprised by one target and one distractor sentence. They subsequently repeated the target sentence. English translations of the sentences: After these squalls there is a terrible chaos (purple), Clouds gather in the blue sky (green). B: Effects of word surprisal and word audibility on word-by-word comprehension accuracy. Surprisal and audibility jointly predicted comprehension accuracy, with a stronger effect of surprisal at low levels of audibility.

To control behavioral performance for peripheral differences in hearing abilities, speech reception thresholds (SRTs) were measured using an adaptive staircase procedure adapted from (Kollmeier et al., 1988; Rysop et al., 2021) including 20 additional stimuli. Distractor intensity was held constant at 70 dB sound pressure level. Target intensity was controlled by a staircase procedure, with a decrease in intensity if participants correctly repeated the full target sentence, and an increase in intensity if participants incorrectly repeated at least one word. Accuracy was rated by the experimenter. Similar as in Rysop et al. (2021), the staircase procedure started with an T/D of 5 dB and a step size of 6 dB. Step size was decreased by a factor of 0.85 following each turning point. SRTs were calculated as the average of the T/D ratios preceding the final 5 turning points. The mean SRT across participants was *M* = −8.16(*SD* = 0.93).

### Stimuli

Stimuli were recorded at 44,100 Hz using a high-quality condenser microphone (Rode NT55) in a sound-proof booth. The distractor sentences started 0.5 seconds prior and lasted for a minimum of 0.3 seconds longer than the target sentences to ensure a complete masking of the target stream. Accordingly, the target sentences were comprised by 5-7 words and lasted for 2-4 seconds. The distractor sentences were comprised by 7-10 words and lasted for 3-5 seconds. The sentence material was retrieved from a German sentence corpus (Schiel & Baumann, 2006) after filtering for the suitable sentence length and excluding sentences that contained names or very unusual words. Additionally, similar suitable sentences were created by GPT-3.5. The prompts used for sentence generation can be retrieved from Table S2. All sentences were loudness normalized to -23 dB LUFS using pyloudnorm (Steinmetz & Reiss, 2021).

### EEG data acquisition

EEG was recorded in a soundproof, electromagnetically shielded booth at a sampling rate of 1,000 Hz using a REFA8 68-channel amplifier system (TMSi, Oldenzaal, the Netherlands), grounded to the sternum. The recording included 63 Ag/AgCl electrodes positioned according to the 10-20 layout in ANT Neuro waveguard original caps as well as two mastoid external mastoid references (A1, A2). Eye movements were recorded using bipolar electrooculogram (EOG) electrodes placed at the outer sides of both eyes and at the top and bottom of the right eye. The EEG signals were monitored throughout the experiment and electrodes showing any visible signs of noise were noted. Impedance was controlled repeatedly during the breaks and electrode preparation was adjusted if deemed necessary by the experimenters.

### EEG preprocessing

EEG preprocessing was conducted using MNE python (Gramfort et al., 2013) and followed recommendations for temporal response function (TRF) analyses (Crosse et al., 2021). The signal was filtered between 0.5 and 15 Hz and re-referenced to the average of the mastoid references. Bad channels were rejected manually based on a larger variance than neighboring channels. Independent component analysis (ICA) with 15 components was conducted to reduce artifacts from blinking and eye movements. The rejected components were selected automatically based on correlations of at least 0.3 with the EOG electrodes. We finally interpolated bad channels using spherical splines (Perrin et al., 1989) and downsampled the signals to 128 Hz for TRF estimation to reduce computation times. The signal was epoched between target sentence onset and offset.

### Features

#### Acoustic features

Acoustic features included stimulus envelopes and acoustic onsets. Additionally, we investigated the neural tracking of word and phoneme onsets. All features were generated separately for target and distractor streams.

Envelopes and acoustic onsets were derived from auditory spectrograms calculated using naplib (Mischler et al., 2023). Spectrogram calculation involved a cochlear filter bank of 128 logarithmically-spaced constant-Q filters and a hair cell model to approximate the human peripheral auditory system. Envelopes were calculated using the mean across all spectrogram channels. Acoustic onsets were derived using the half-wave rectified derivative of the envelope (Brodbeck et al., 2018).

To derive word and phoneme onsets, stimuli and text transcripts were automatically aligned using forced alignment available in the WebMAUS Basic module of the BAS Web Services (Kisler et al., 2017; Schiel, 1999). Naturally, the word and phoneme onset predictors were highly correlated, as every word onset coincides with a phoneme onset. However, the inclusion of both predictors was necessary due to the need to control the higher-level analyses for possibly distinct onset profiles in words and phonemes. To avoid artifacts at sentence onset, the first word and phoneme onsets in each sentence were modeled as a separate predictor.

### Linguistic features

To quantify the neural tracking of linguistic features, we investigated features at the word and phoneme levels, as previous studies showed that these features are significantly tracked in continuous listening paradigms (Brodbeck et al., 2018; Gillis et al., 2021).

### Word level features

Word frequency was derived from the SUBTLEX-DE subtitles corpus (Brysbaert et al., 2011). Word frequency was defined as the logarithm of word occurence per million, as provided in the SUBTLEX-DE database.

Word surprisal is defined as the inverse probability of each word given the preceding context. Surprisal of word i is defined as the negative logarithm of word probability:

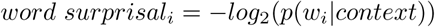

with (*p*(*w_i_*|*context*) being the word probability given the preceding sentence context derived from GPT-2 trained on a German text corpus (Schweter, 2020). Since we used isolated sentences, surprisal estimates of the first words in each sentence cannot be interpreted based on prior context. For the distractor sentences, an average of two word onsets were omitted by cutting the first 500 ms before target sentence onset. To ensure compatibility for the target sentences and to provide sufficient context for the surprisal calculation, we omitted the first two words in each target sentence from the higher-level analyses (Slaats et al., 2023).

### Phoneme level features

Phoneme surprisal reflects probability of each phoneme given the preceding phonemes in the current word. It is based on a probability prior captured by word frequency. Phoneme surprisal was defined as the inverse conditional probability of each phoneme, given the preceding phonemes in the same word:

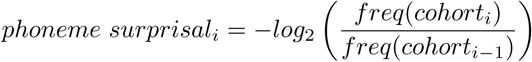

*Cohort_i_* is the activated cohort of words given the current phoneme. This is the amount of words in the vocabulary that could be formed from the given sequence of phonemes.

*Freq*(*cohort*) is the sum of the word frequencies of all words in the cohort.

Cohort entropy reflects the uncertainty about the next phoneme. It is therefore defined as the Shannon entropy of the activated cohort (Brodbeck et al., 2018). Entropy at cohort i is defined by:

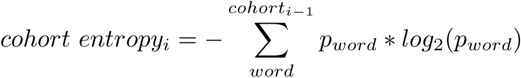

with *p_word_* being the probability of each word as reflected by its relative frequency. To appropriately separate the word and phoneme levels, we only used phoneme events that did not correspond to word onsets, i.e., omitting the first phonemes in each word. All features were min-max normalized to the range from 0 to 1 to avoid negative values before entering the TRF.

### Speaker characteristics

As mentioned above, target and distractor sentences were presented by a male and a female voice, respectively. This was done to prevent confounding effects of the speaker, potentially dampening the effects. To control for the differences introduced by the speakers, the analysis was controlled for fundamental frequency. Fundamental frequency was calculated using the pYIN algorithm (De Cheveigné & Kawahara, 2002; Mauch & Dixon, 2014) available in the python package librosa (McFee et al., 2024).

Additionally, we controlled the linguistic analyses for a set of variables capturing perception of speaker-specific characteristics. To obtain these speaker characteristics, a subsample of the participants (N = 23) took part in an online experiment after completing the EEG experiment. Here, participants were presented with ten sentences spoken by the same female and male speakers used in the EEG experiment. After each sentence, they were asked to rate the speakers on a set of 7 characteristics: age, educatedness, dominance, attractiveness, health, and professionalism (Lavan et al., 2024). The characteristics were rated on 8 point ratings scales and the ratings were summarized across trials. Using two-sided paired Wilcoxon tests corrected for multiple comparisons using false discovery rate (FDR), we revealed significant differences between the speakers on 3 of the characteristics (age [W = 21.5, p = .001], attractiveness [W = 8.5, p = .001], and dominance [W = 56, p = .031]). The speaker-specific mean ratings of these characteristics were added into the acoustic model containing the respective characteristic at each word onset in order to control the linguistic analysis for the speaker characteristics.

### Temporal response function modeling

In order to investigate how comprehension performance is correlated with the interplay of acoustic and linguistic features of the attended and unattended streams, we used multivariate TRFs to quantify how features were neurally tracked. The model fits were then entered into a generalized linear mixed effects model to access the effect of feature tracking on comprehension.

To investigate the neural encoding of acoustic and linguistic features, we used multivariate TRFs. The analysis was implemented in mTRFpy (Bialas et al., 2023) using multiple linear regression with Tikhonov regularization. The TRF is a kernel that describes the linear transformation from an ongoing stimulus to an ongoing neural response for a specified set of time lags. When applying forward modeling, the TRF is used to predict the EEG signal from a set of stimulus features. To obtain the predicted EEG signal, the TRF is convolved

with the speech features of the target and distractor sentences. The prediction performance is then obtained by measuring the correlation between the predicted EEG signal and the actual signal. The prediction performance can then be interpreted as the amount of neural tracking of the speech features. Feature representations were investigated in time lags from -100 to 800 ms relative to event onset.

For the TRF analysis, we divided the trials into correct (i.e., all words of the sentence were repeated correctly) and incorrect trials. The criterion for correct sentences was deliberately chosen to be strict in order to ensure that participants grasped the full sentence context. Separate models for correct and incorrect trials were trained across participants, as previously recommended for small datasets (Crosse et al., 2021; Jessen et al., 2021). This was done to increase the amount of training data, as the reliability of TRF analyses has been shown to benefit from an increase in training data, especially for sparse linguistic features (Mesik & Wojtczak, 2023). Additionally, we aimed to reduce the effect of varying numbers of correct and incorrect trials across participants. We used an 80-20 cross-validation with 50 randomly sampled folds following recent recommendations for evaluating brain decoding models (Varoquaux et al., 2017). For each fold, 8 participants were randomly selected to comprise the test set. Data from these participants was not used for the model training. Models for correct trials were, on average, trained on 3831 trials, models for incorrect trials were trained on a mean number of 3169 trials. TRFs were evaluated for the duration of the target sentences, omitting the initial and the terminal segments where only the distractor sentence was presented.

We investigated two separate models: an acoustic and a linguistic TRF model. In the acoustic model, we included the lower-level acoustic features (i.e., envelope and acoustic onsets) as well as word and phoneme onsets. The linguistic model included all linguistic features on the word and phoneme levels. Naturally, higher-level features are highly correlated to lower-level features in speech, making it important to control analyses investigating higher-level features for lower-level variance (Gillis et al., 2021). We therefore controlled the higher-level analyses for the acoustic features. For this control, we conducted a separate TRF analysis predicting the EEG signals from all acoustic variables and word and phoneme onsets, including the speaker characteristics described above, with no cross-validation. Model predictions from this analysis were then subtracted from the original EEG data to generate residualized EEG responses that do not contain responses resulting from acoustic features. All analyses involving linguistic features were conducted on these residualized EEG responses. To isolate the influence of the distinct feature groups (see Table 1), we systematically shuffled the features of interest (see below). The regularization parameter was fitted separately for acoustic and linguistic models, but it was commonly fitted across participants and correct and incorrect model variants to avoid overfitting and ensure comparability between models. Acoustic models were fitted using a regularization parameter of 0.1 and linguistic models were fitted with a regularization of 1e-7.

**Table 1:**
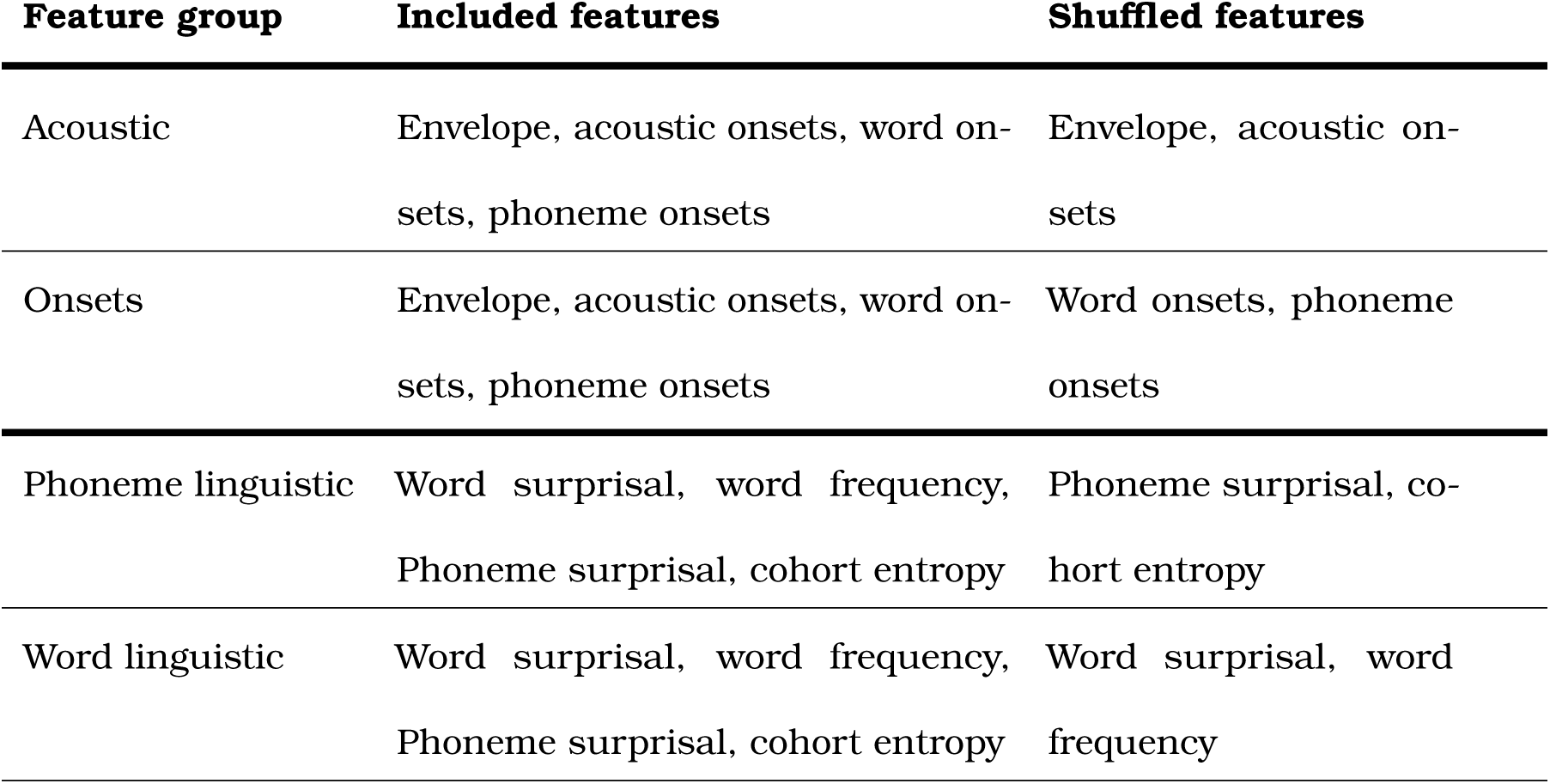
TRF Models and Corresponding Speech features.

### Statistical Inference

We used a shuffling approach to infer statistical significance of the TRF model fits. This was done to make sure that all models were trained on the same number of variables and improvements in model performance cannot solely be explained by the addition of variables. We grouped the individual features into four groups to avoid multicollinearity issues in regression model fitting. The feature groups and their corresponding features are shown in Table 1. We conducted 50 shuffling iterations for each model variant. In each iteration, we systematically shuffled the features of interest while keeping all other input features and model parameters identical. We used different shuffling methods for acoustic and linguistic models, corresponding to the characteristics of the respective features. For the acoustic models, trial labels were shuffled. For the linguistic models, we employed a more conservative approach of shuffling the values for each feature while preserving the onset times. Feature values were shuffled globally across sentences and separately for each feature to retain distributional properties. Due to the sparse nature of the linguistic regressors, we constrained the model evaluation to all time points from 0 to 800 ms after an event onset, given that these are the latencies at which the events can realistically influence the neural response.

Statistical significance was inferred by subtracting the mean shuffled model fits from the unshuffled model fits. Statistical inference was performed using mass-univariate two tailed one sample t-tests with threshold-free cluster enhancement and a cluster forming threshold of *α* = 0.05. This analysis yields p-values corrected for multiple comparisons (Maris & Oostenveld, 2007).

### ROI selection

To select electrodes of interest, a separate TRF analysis was conducted on the trials of the control condition. This condition was comprised by 40 trials per participant, and the data from these trials were left out from the main analysis to avoid double dipping. The analysis was conducted using the same model variants, features, and cross-validation procedure as described above for the main analysis. Additionally, the shuffling procedure as well as the residualization was applied in the same way as described above for the main analysis. Based on previous studies, we selected three clusters that can be assumed to play important roles in acoustic and linguistic processing (Gillis, Kries, et al., 2023). As in Gillis, Kries, et al. (2023), we used temporal and centro-parietal electrode selections, as these can be assumed to be involved in lower- and higher-level speech processing. We additionally selected a fronto-central cluster, as acoustic tracking is commonly shown to be strongest in the corresponding sensors (Gillis, Kries, et al., 2023; Gillis, Vanthornhout, & Francart, 2023; Lesenfants et al., 2019). Electrodes of interest were selected by choosing the cluster with the highest model fit in the control condition. Topographies for the control condition along with the selected clusters can be retrieved from Figure S2.

To test for differences between correct and incorrect trials in the selected clusters, we conducted paired, two-sided t-tests on the mean model fits per feature. The p-values resulting from these comparisons were corrected for multiple comparisons using FDR. To allow comparisons between these t-tests and the mixed model analysis independent of the number of observations, we provide effect sizes along with the statistical significance estimates.

### Neuro-Behavioral Correlations

The speech recordings were automatically transcribed offline using whisper large-v2 (Bain et al., 2023). Subsequently, the transcriptions were manually controlled. We then assessed the overlap between the transcribed response and the target sentence, marking each word as accurately repeated or inaccurate.

Word-by-word repetition performance was predicted from word surprisal and word audibility. The amount of masking varies on a moment-by-moment basis in competing speech stimuli, leading to some speech segments being relatively well audible. This has been described as glimpsing (Cooke, 2006). Glimpsed segments can inform the comprehension of segments that coincide with higher amounts of acoustic masking. To estimate the amount of available glimpses in each target word, we calculated the so-called “glimpse rate” for each target word segmentation (Cooke, 2006). This means that we computed the available amount of acoustic information in each word by comparing target and distractor spectrogram amplitudes. The glimpse rate was defined as the proportion of spectrotemporal events that exceed a certain local T/D threshold within each word. This threshold was set to -5 dB based on previous work (Cooke, 2006).

To estimate the effects of word surprisal, audibility, and their interaction on word repetition performance (correct versus incorrect word repetition) in the experimental conditions, we calculated a generalized linear mixed model with a logistic link function using lme4 in R (Bates, 2014). P-values were corrected for multiple comparisons using FDR. The analysis was controlled for word frequency, sentence length (in words), word order, T/D condition, and accuracy in the control condition. We included random intercepts for subjects and sentences. To investigate the relationship between TRF model fits and behavioral performance, standardized trial-wise model fits for all model variants were entered into the logistic regression model.

To rule out issues of multicollinearity, we calculated variance inflation factors using the R package car (Fox & Weisberg, 2019). The variance inflation factors ranged from 1.00 to 2.26 and we therefore assume no issues of multicollinearity (O’brien, 2007). We additionally inspected the correlations between the model fits and revealed that none of the model fits were strongly correlated. All correlations are displayed in Figure S4. The full model explained 15.9 % of variance in comprehension considering fixed effects only, and 50.8 % of variance if random effects were taken into account.

We conducted a supplementary analysis to further investigate the interactions between correct and incorrect trials across the feature groups and the streams. Here, we computed a linear mixed model predicting the standardized TRF model fit from condition (correct versus incorrect), feature group (acoustic versus onsets versus phoneme linguistic versus word linguistic), stream (target versus distractor) and their interactions. Due to heteroskedasticity,

we calculated a generalized linear mixed model accounting for different variances of the conditions using the R package glmmTMB (Brooks et al., 2017; McGillycuddy et al., 2025). The models included random intercepts for participants. Post-hoc comparisons were conducted using the R package emmeans (Lenth, 2024).

### Audibility control analysis

Due to the competing speech paradigm, speech clarity varies on a word-by-word basis, influencing comprehension. To infer influences of speech clarity on lower-level processing, we calculated separate TRF model fits based on the word audibility of each target word for both conditions (correct and incorrect). Separate model fits were obtained by dividing the observed and the predicted EEG signals into segments of low and high audibility. Predicted EEG signals resulted from the cross-validated predictions of the TRF models trained on all data. Then, the correlation between both signals was measured separately for segments of high and low audibility. The division of the words was done separately for correct and incorrect trials, so that each split would include half of the signal in each condition (Tezcan et al., 2023). Model fits were summarized within the previously defined clusters with maximal model fit (see above). Word audibility is influenced by word-by-word speech clarity, as well as by the T/D condition of the sentences. To combine word-by-word audibility and sentence T/D, we summed the standardized glimpse rate of each target word segmentation (Cooke, 2006) and the T/D condition of the respective sentence. To infer statistical significance, we conducted paired samples t-tests comparing model fits for segments with high and low audibility separately for correct and incorrect trials. Due to the explorative nature of the analysis, we report uncorrected as well as FDR corrected p-values.

### Code Accessibility

Custom code for all analyses will be made available upon acceptance and publication of the manuscript.

## Results

### Behavioral Results

#### Comprehension is modulated by linguistic features particularly at low audibility

In a generalized linear mixed model, we predicted word-by-word comprehension performance from word-level audibility and surprisal. Word-level audibility was derived from the proportion of the target signal exceeding a certain local SNR threshold (e.g., the “glimpse rate”). Results revealed that audibility, surprisal, as well as their interaction predicted word repetition performance. Figure 1 displays the interaction between word audibility and surprisal, indicating that the effect of surprisal was stronger at low levels of audibility (see Table 2 for the full set of results). This means that more predictable words were easier to understand than less predictable words when it was more difficult to hear the attended speech stream.

**Table 2:** Generalized Linear Mixed Model outputs predicting comprehension performance.

|  | Estimate | Std. Error | CI | z | p |
| --- | --- | --- | --- | --- | --- |
| Audibility | 0.39 | 0.02 | [0.35, 0.43] | 19.64 | <.001** |
| Surprisal | -0.39 | 0.03 | [-0.44, -0.33] | -14.93 | <.001** |
| Audibility:Surprisal | 0.12 | 0.02 | [0.08, 0.16] | 5.57 | <.001** |
| Acoustic target | -0.03 | 0.02 | [-0.06, 0.01] | -1.54 | .149 |
| Onsets target | 0.07 | 0.02 | [0.04, 0.11] | 4.19 | <.001** |
| Word linguistic target | 0.06 | 0.02 | [0.03, 0.1] | 3.84 | <.001** |
| Phoneme linguistic target | 0.03 | 0.02 | [-0.01, 0.06] | 1.57 | .149 |
| Acoustic distractor | 0.05 | 0.02 | [0.02, 0.09] | 3.18 | .003** |
| Onsets distractor | -0.02 | 0.02 | [-0.05, 0.01] | -1.26 | .232 |
| Word linguistic distractor | -0.04 | 0.01 | [-0.07, -0.01] | -2.6 | .014* |
| Phoneme linguistic distractor | 0.00 | 0.02 | [-0.03, 0.03] | -0.27 | .790 |
| Target-to-Distractor ratio (dB) | 0.49 | 0.01 | [0.47, 0.51] | 41.09 | <.001** |
| Word frequency | -0.07 | 0.02 | [-0.12, -0.02] | -2.85 | .007** |
| Word order | -0.07 | 0.01 | [-0.1, -0.04] | -4.62 | <.001** |
| Sentence length | -0.15 | 0.12 | [-0.39, 0.09] | -1.22 | .236 |
| Control accuracy | 0.21 | 0.1 | [0.02, 0.4] | 2.21 | .038* |
| Speech reception threshold | 0.28 | 0.05 | [0.18, 0.38] | 5.38 | <.001** |

\*\* p < .01, \* p < .05

### Temporal response function analysis

#### Acoustic features are significantly tracked for target and distractor sentences

To determine regions with significant model fits, we conducted mass-univariate two tailed one sample t-tests on the difference between TRF model fits and shuffled model fits. Acoustic features were significantly tracked in target and distractor sentences. As displayed in Figure 2 A, the model fit was strongest in frontal and central electrodes. Figure S3 shows the full topographies and results from the mass-univariate statistics. Paired t-tests revealed a significant difference between TRF model fits for correct and incorrect trials for the distractor (*t* = 4.91*, df* = 42*, p_corrected_ < .*001*, d* = 0.75), but not for the target stream (*t* = 0.26*, df* = 42*, p_corrected_* = .80*, d* = 0.04). This indicates that the TRF model fits for the distractor acoustics were higher in correct than in incorrect trials.

**Figure 2:**
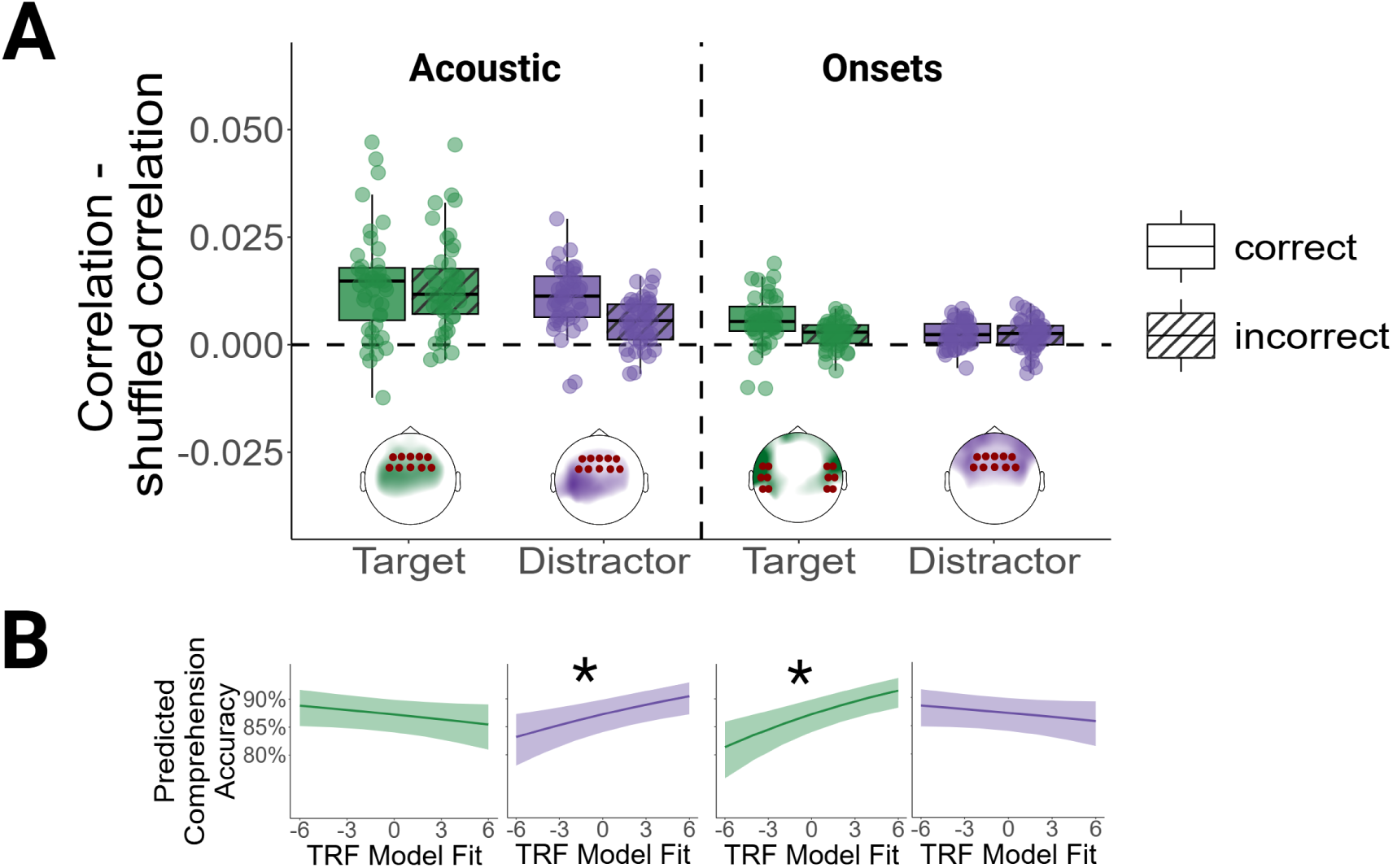
Acoustic TRF model fits, clusters, and model predictions. A: Fits for the acoustic features. Each dot represents the fit for one participant. Correlations indicate the correlations between the EEG signals predicted by the temporal response functions and the actual EEG signals. The topographies display the spatial organization of the model fits, with colors indicating the magnitude of the model fit. The model fits were z-scored within conditions to facilitate the visual inspection of their spatial organization. Electrodes belonging to the selected clusters are marked by red circles. B: Generalized linear mixed model predictions for the acoustic features. Correct and incorrect trials were combined in the generalized linear mixed effects model. Acoustic refers to the model fits for the envelope and the acoustic onsets features. Onsets refers to the model fits for word and phoneme onsets. * p < .05

Similarly, word and phoneme onsets were significantly tracked for target and distractor sentences. For the target sentences, maximal model fits were observed in temporal electrodes. For the distractor sentences, the strongest fits were observed in frontal electrodes. The subsequent analyses of model fits and model weights were constrained to clusters showing the best prediction performance in a held-out control condition. These clusters are displayed in Figure 2 A. Paired t-tests revealed a significant difference between TRF model fits for correct and incorrect trials for the target stream (*t* = 3.63*, df* = 42*, p_corrected_* = .003*, d* = 0.55), indicating that the model fits were higher in correct than in incorrect trials. There was no significant difference between correct and incorrect trials for the distractor stream (*t* = 0.26*, df* = 42*, p_corrected_* = .80*, d* = 0.04).

### Linguistic features are significantly tracked for target sentences

Mass-univariate statistics revealed significant clusters for word level linguistic features in the target sentences for correct and incorrect trials. As shown in Figure 3 A, prediction performance peaked in central posterior electrodes for the word level linguistic features. A paired t-test did not reveal a significant difference between TRF model fits for correct and incorrect trials (*t* = 1.61*, df* = 42*, p_corrected_* = .18*, d* = 0.25). Phoneme level linguistic features were significantly tracked for the target sentences in correct trials, with the highest prediction performance in temporal and frontal sensors as shown in Figure 3 A. Phoneme level linguistic features were not significantly tracked for the target sentences in incorrect trials. A paired t-test did not reveal a significant difference between TRF model fits for correct and incorrect trials (*t* = 1.26*, df* = 42*, p_corrected_* = .29*, d* = 0.19).

**Figure 3:**
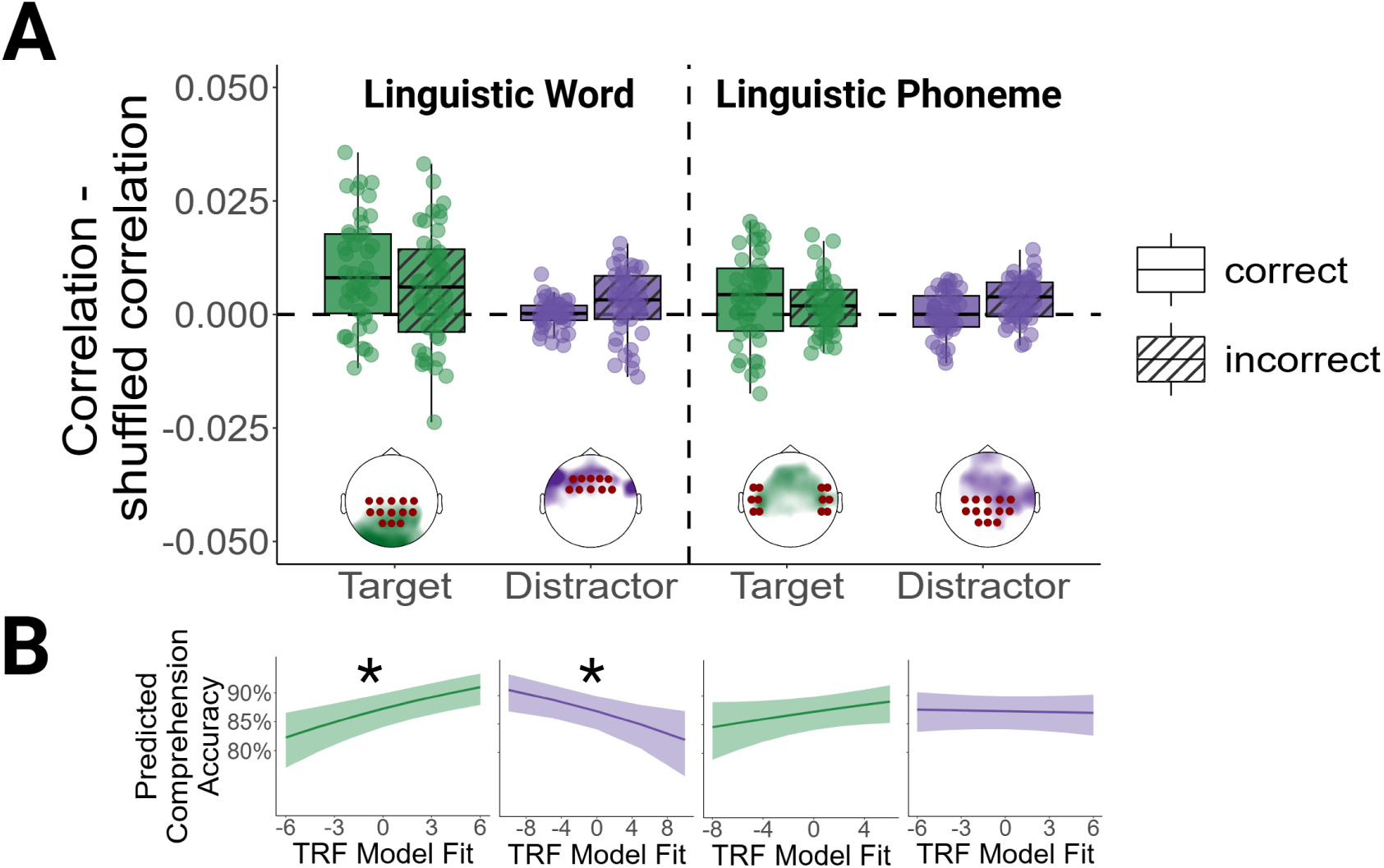
Linguistic TRF model fits, clusters, and model predictions. A: Fits for the linguistic features. Each dot represents the fit for one participant. Correlations indicate the correlations between the EEG signals predicted by the temporal response functions and the actual EEG signals. The topographies display the spatial organization of the model fits, with colors indicating the magnitude of the model fit. The model fits were z-scored within conditions to facilitate the visual inspection of their spatial organization. Electrodes belonging to the selected clusters are marked by red circles. B: Generalized linear mixed model predictions for the acoustic features. Correct and incorrect trials were combined in the generalized linear mixed effects model. Linguistic word model fits refer to the model fits for word surprisal and word frequency. Linguistic phoneme model fits refer to the model fits for phoneme surprisal and cohort entropy. * p < .05

### Linguistic features are only tracked for distractor sentences in incorrect trials

As shown in Figure S3, word and phoneme level linguistic features for unattended distractor sentences were only significantly tracked in incorrect trials. This means that no significant tracking of higher-level features in the unattended speech stream emerged if comprehension of the target stream was intact. In contrast, higher-level features for the unattended stream were neurally tracked when comprehension of the attended stream collapsed. Mass-univariate statistics revealed significant prediction performance for word level linguistic features in frontal sensors in incorrect trials. Model fits for linguistic phoneme features were significant in frontal and central sensors. A paired t-test revealed a significant difference between TRF model fits for correct and incorrect trials for the phoneme features (*t* = −2.82*, df* = 42*, p_corrected_* = .02*, d* = −0.43). This difference indicates that the model fits were higher for the phoneme distractor features in incorrect than in correct trials. There was no significant difference between the model fits for correct and incorrect trials for the word level features (*t* = −2.02*, df* = 42*, p_corrected_* = .09*, d* = −0.31).

### Neuro-Behavioral Correlations

#### Enhanced tracking of the target stream supports comprehension

To assess effects of the neural tracking of lower-level and higher-level features on comprehension, we predicted word-by-word comprehension accuracy from the trial-wise TRF model fits. We summarized model fits within the clusters displayed in Figures 2 A and 3 A. Focusing on the target stream, comprehension accuracy was positively correlated with the neural tracking of target onsets and word level linguistic features. This means that the neural tracking of target onsets and target word level information was stronger when the sentences were comprehended. Model parameters can be retrieved from the second section of Table 2. Model predictions are visualized in Figures 2 B and 3 B.

As a result of training the TRF models across participants, the model weights do not reflect subject-specific, independent observations, which complicates statistical inference. Nevertheless, descriptive analysis of the weights shown in Figure 4 revealed similar results. Weights for the word linguistic features in correct trials were characterized by stronger responses starting from 400 ms. This corresponds to the typically observed time range of linguistic word level effects (Hsin et al., 2023; Kutas & Federmeier, 2011; Lau et al., 2008; Van Petten & Luka, 2012), which have been shown to be delayed in noise and degraded speech (Connolly et al., 1992; Obleser & Kotz, 2011; Strauß et al., 2013). In contrast, weights for incorrect trials showed smaller amplitudes in late windows. Weights for the onsets revealed stronger early responses in correct trials than in incorrect trials. Fits for the phoneme linguistic features only reached significance in incorrect trials for the distractor sentences and in correct trials for the target sentences.

**Figure 4:**
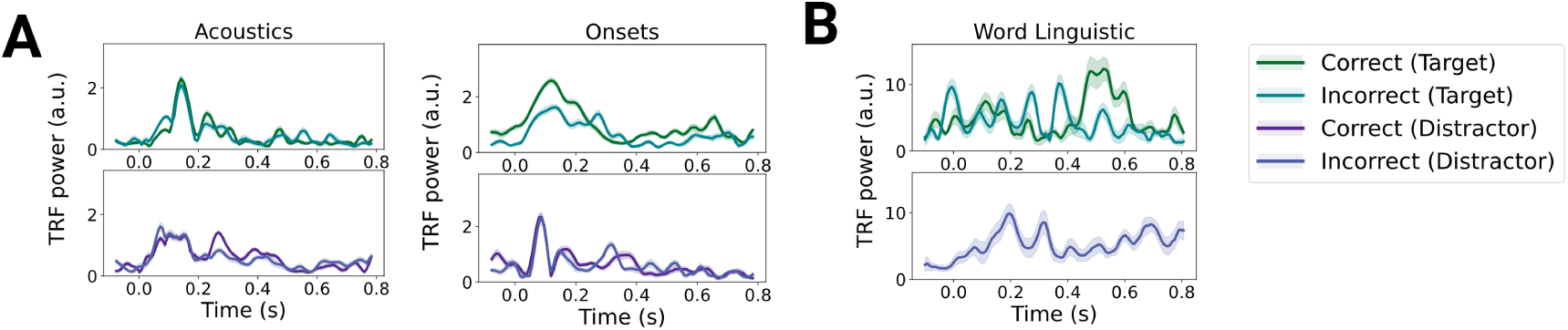
TRF weights. A: TRF weights for the acoustic models. B: TRF weights for the linguistic word level model. Fits for the linguistic features did not reach significance for the distractor in correct trials and were therefore omitted.

### Distractor processing is beneficial at lower and detrimental at higher levels

By predicting target comprehension accuracy from the trial-wise TRF model fits for the distractor stream, we revealed that acoustic tracking of the distractor was positively correlated with comprehension performance. In contrast, word level distractor tracking was negatively correlated with comprehension performance. Full results can be retrieved from Table 2. This shows that, only at higher levels, distractor processing showed a distracting effect, i.e., performance for target comprehension was negatively impacted when distractor information was overrepresented.

A supplementary analysis applying a linear mixed model predicting TRF model fits from accuracy, feature group, and stream (target versus distractor) revealed results consistent with our main analysis. The model revealed a significant three-way interaction between accuracy, feature group, and stream. Post-hoc comparisons revealed that the interaction between feature group and accuracy only reached significance for the distractor stream (F = 10.23, p < .001), but not in the target stream (F = 0.57, p = .637). This means that, only in the distractor stream, there was a significantly different effect for the higher-level features when compared with the lower-level acoustic features. The full results for the linear mixed model can be retrieved from Table S1 and Figure S5. These results confirm the results from the main analysis by revealing that the effect of accuracy on distractor representation follows different patterns in lower and higher representational levels.

As mentioned above, interpretation of the TRF model weights is difficult given the low variance across iterations. However, we observed that distractor weights at the word level for incorrect trials were primarily characterized by early responses, as shown in figure 4 B. This differs from the time scale of the strongest effects in the target streams.

### Influences of Audibility on TRF Model Fits

#### Lower level processing is influenced by speech clarity

To infer bottom-up acoustic influences on the encoding of different features, we calculated separate model fits in sections with high and low word audibility. We then conducted paired samples t-tests comparing high and low audibility model fits seperately for incorrect and correct trials. Significant results can be visually inspected in Figure 5. Results revealed that target acoustic tracking was higher in segments of high audibility compared to segments of low audibility in incorrect trials (*t* = 3.27*, df* = 42*, p_corrected_* = .026) and in correct trials (*t* = 2.83*, df* = 42*, p_corrected_* = .043).

**Figure 5:**
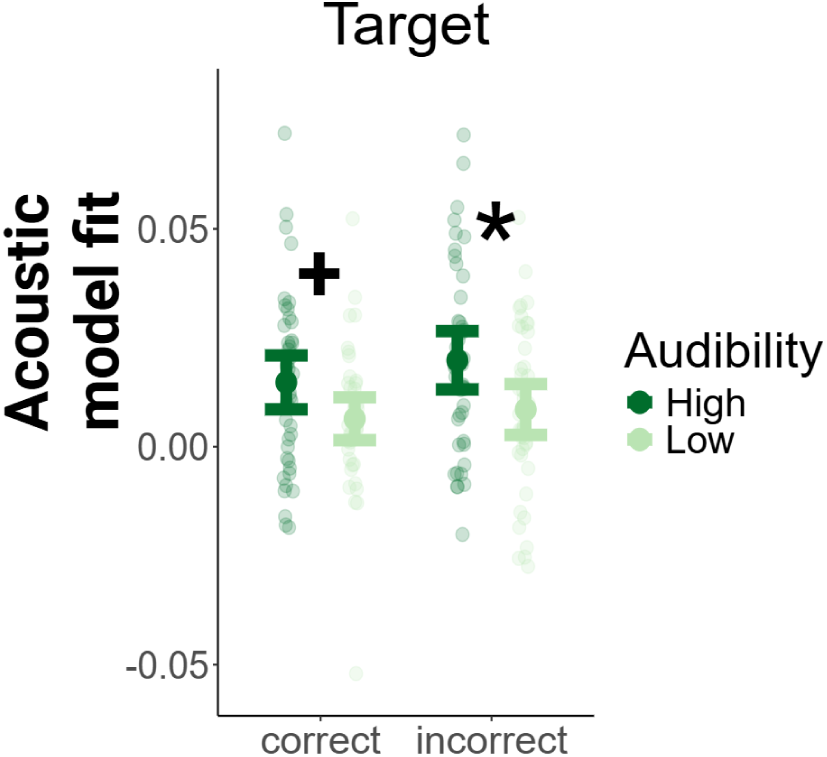
Acoustic TRF model fits by Audibility. Increased acoustic target tracking in high audibility. Points show the means and their 95% confidence intervals. Transparent dots show the raw individual subject data. * p < .05 (FDR corrected for multiple comparisons), + p < .05 (uncorrected for multiple comparisons)

## Discussion

In this study, we examined how acoustic and linguistic processing contribute to speech comprehension within a competing speech paradigm. As a main finding, we show that the attentional engagement with the target stream positively contributed to comprehension at lower and higher hierarchical levels. In contrast, the neural tracking of distractor information was positively correlated with comprehension at lower levels, whereas higher level distractor engagement had a negative effect on comprehension. These results suggest that target enhancement and distractor suppression jointly support comprehension. However, increased attention to distractor features only impedes comprehension at higher hierarchical levels, but not at lower acoustic processing levels.

### Acoustic and linguistic features influence comprehension

On the behavioral level, we showed that word predictability and word audibility jointly and interactively predicted comprehension performance. The effect of word predictability was strongest when audibility was low and the target words were therefore more difficult to hear. This is consistent with previous literature showing influences of word predictability on comprehension performance (Kalikow et al., 1977), as well as on behavioral indicators of processing demands, such as reading times (Smith & Levy, 2013).

To obtain a fine-grained behavioral comprehension measure, we adopted the widely used method of sentence repetition (Kalikow et al., 1977; Rysop et al., 2021; Stickney & Assmann, 2001). To infer the effect of working memory constraints on sentence repetition, we assessed task-specific working memory performance in a control condition comprised by sentences that were well comprehended by the participants. The analysis was controlled for the accuracy in this control condition. Accuracy in the control condition amounted to an almost perfect accuracy of 97% of words across participants. Thus, we assume that comprehension scores in the remaining conditions truly reflect comprehension instead of working memory constraints.

### Target enhancement predicts comprehension

Consistent with previous studies (Etard & Reichenbach, 2019; Iotzov & Parra, 2019; Lesenfants et al., 2019; Tune et al., 2021; Vanthornhout et al., 2018), we found that the stronger neural representation of the target stream positively predicted comprehension performance. This effect was present for word and phoneme onsets, suggesting that successful word and phoneme segmentation is crucial for comprehension. This is congruent with a recent study revealing that the neural representation of word onsets differentiates between intelligible and unintelligible speech while controlling for acoustic differences (Karunathilake et al., 2023). Additionally, the neural representation of word level linguistic information was positively predictive of comprehension performance. This effect is unsurprising given the previous literature (Broderick et al., 2018; Holcomb, 1993; Karunathilake et al., 2023; Obleser & Kotz, 2011). However, extending previous work, we provide direct evidence that increased attention towards lower- and higher-level neural representations of the attended stream is a neural indicator of comprehension success in a challenging multitalker situation. Importantly, higher level tracking was predictive of comprehension performance while controlling for lower level acoustic neural tracking, underlining its importance for the prediction of successful language comprehension.

Interestingly, the neural representation of acoustic target information (i.e., the envelope and acoustic onsets) was not predictive of comprehension performance. This seems to contrast previous studies demonstrating that acoustic tracking is sensitive to speech clarity (Ahissar et al., 2001; Doelling et al., 2014; Etard & Reichenbach, 2019; Iotzov & Parra, 2019; Lesenfants et al., 2019; Vanthornhout et al., 2018). In our audibility control analysis, we show that target envelope tracking was indeed sensitive to word-by-word speech clarity. However, we did not find an effect of target envelope tracking on speech comprehension. Similarly, several previous studies showed that acoustic tracking is not directly related to speech comprehension (Kösem et al., 2023; Verschueren et al., 2022). This is consistent with a recent study showing that higher level representations (i.e., word onsets and word level linguistic features) differentiate intelligible from unintelligible speech, while controlling for speech acoustics (Karunathilake et al., 2023). Collectively, the previous and present findings suggest that the neural tracking of higher-level linguistic information is crucial for higher-level comprehension while lower-level acoustic target tracking is related to speech clarity.

### Distractor attention is differentially related to comprehension

In contrast to the representation of target information, previous findings on the neural representation of distractor information have been more heterogeneous. By disentangling acoustic and linguistic processing levels, we show that lower level distractor representation does not interfere with comprehension of the target speech stream. In fact, acoustic tracking of the distractor stream was positively predictive of comprehension, suggesting that stronger distractor tracking at the acoustic level may facilitate comprehension. This may be attributed to the dominance of the distractor in the acoustic speech mixture, with stronger distractor tracking reflecting greater overall listening engagement. Additionally, improved acoustic tracking of the distractor stream may support comprehension by aiding stream segregation (Fiedler et al., 2019; Rimmele et al., 2012). Consistent with previous studies, we show that the attentional competition between attended and unattended signals does not primarily operate at the acoustic level (Golumbic et al., 2013; Orf et al., 2023), as acoustic tracking persists even for the unattended speech stream (Ding & Simon, 2012; Horton et al., 2014; Kaufman & Golumbic, 2023; O’sullivan et al., 2015).

In contrast, higher level distractor tracking was negatively correlated with comprehension, providing direct evidence for a distraction effect. Thus, the presence of higher-level distractor tracking indicates that more attention was allocated to the unattended stream, leading to lower comprehension of the attended stream. Only higher level distractor representation interfered with target comprehension, indicating that, only at this higher level, distractor disengagement becomes perceptually adaptive. Additionally, we replicate the finding that higher level tracking typically does not occur for unattended speech (Brodbeck et al., 2018; Broderick et al., 2018), as long as listeners successfully allocate selective attention. We did not reveal any significant representation of word or phoneme level linguistic features for the distractor sentences in correct trials. Crucially, these representations were present for incorrect trials, suggesting that significant neural tracking of higher level distractor information only emerged when attention allocation to the target stream failed. Interestingly, higher level representations of the distractor sentences were present in different sensors and latencies than higher level representations of the target sentences. Word level linguistic representations of correctly reproduced target sentences were characterized by late responses in central-posterior sensors, which is consistent with previous results using temporal response functions (Gillis et al., 2021) and traditional evidence in event-related potentials, including the N400 as well as later positive components (Aurnhammer et al., 2021; Hsin et al., 2023; Kutas & Federmeier, 2011; Lau et al., 2008; Šoškić et al., 2022; Van Petten & Luka, 2012). Thus, this response profile seems to reflect successful linguistic processing of the target sentences. In contrast, the highest prediction performance for distractor word level features was located in frontal sensors. Additionally, the weights for distractor word level features showed earlier peaks than the weights for target word level features. This response profile does not correspond to typically observed signatures of linguistic processing. Instead, the responses may reflect suppression processes in response to higher level distractor information. The potentially different mechanisms in response to target and distractor linguistic features should be further uncovered in future investigations.

## Conclusion

This study provides evidence that target enhancement as well as distractor suppression at different hierarchical levels are predictive of comprehension performance. While target processing at lower and higher hierarchical levels were positively correlated with compre-hension, distractor influences were positively correlated at lower and negatively correlated at higher levels. This underlines the relevance of the distinct roles of acoustic and linguistic representations in attended and unattended streams when attempting to unravel the neural mechanisms contributing to selective attention and distraction in multitalker environments.

## Acknowledgments

We thank Heike Boethel for her support in data acquisition.

