## supplementary material for "Attentional engagement with target and distractor streams predicts speech comprehension in multitalker environments"

Figure S1: Audiometric measurements

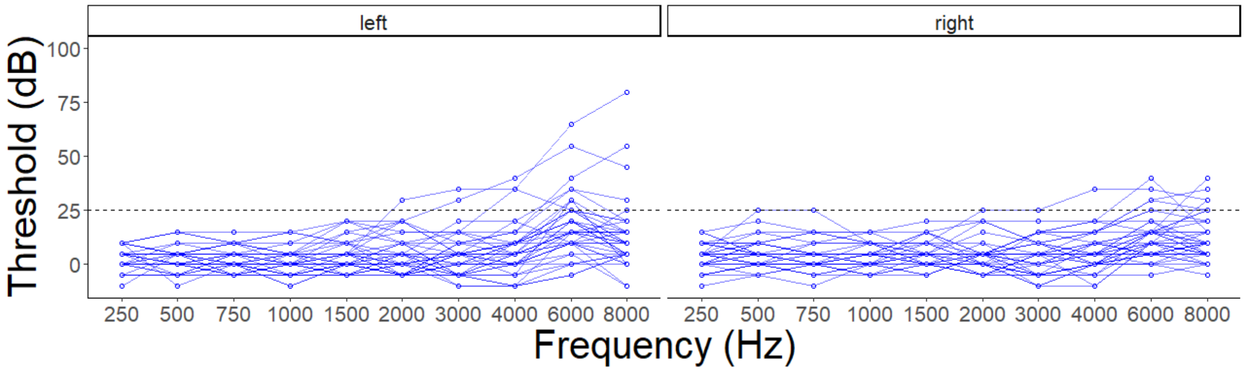


Audiometric thresholds for all participants. Audiometry was conducted prior to starting the experiment. All participants had mean hearing thresholds below 25 dB in at least one ear.

Figure S2: Topographies in the control condition

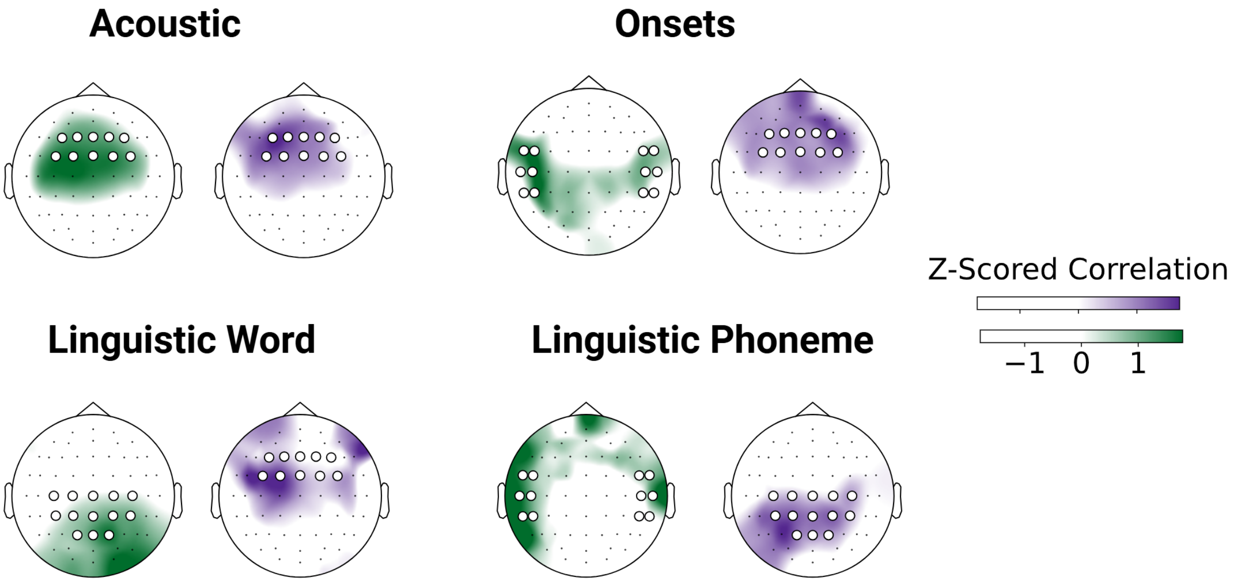


Z-Scored model fits in the control condition used to select the regions of interest for the main analysis. Control trials were held out from the main analysis to avoid double dipping.

Figure S3: Topographies and significant electrodes

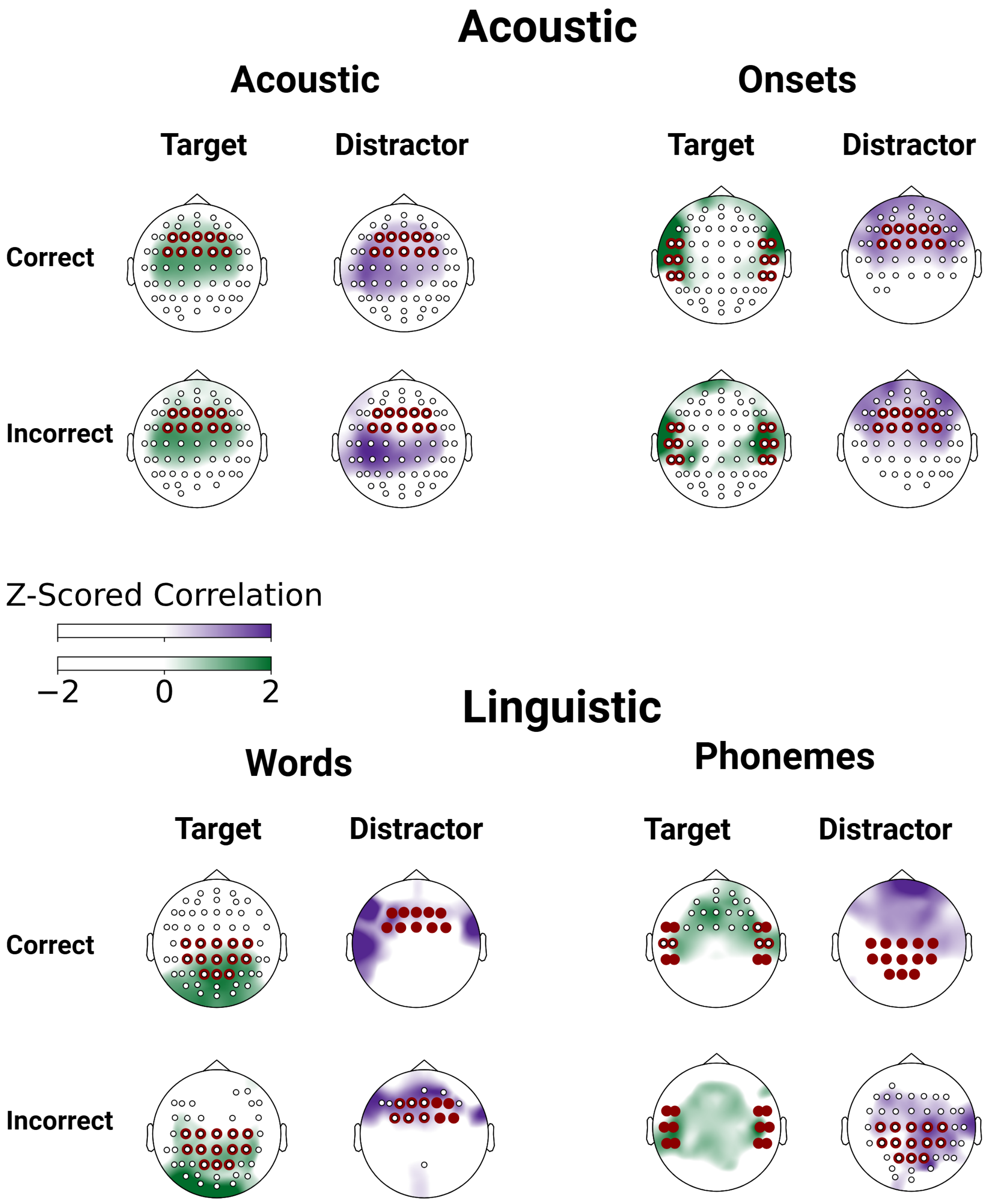


Significant electrodes from zero in the mass-univariate two-sided t-tests are highlighted by white circles. Electrodes used in the analyses are highlighted by red circles.

Figure S4: Correlations between the model fits

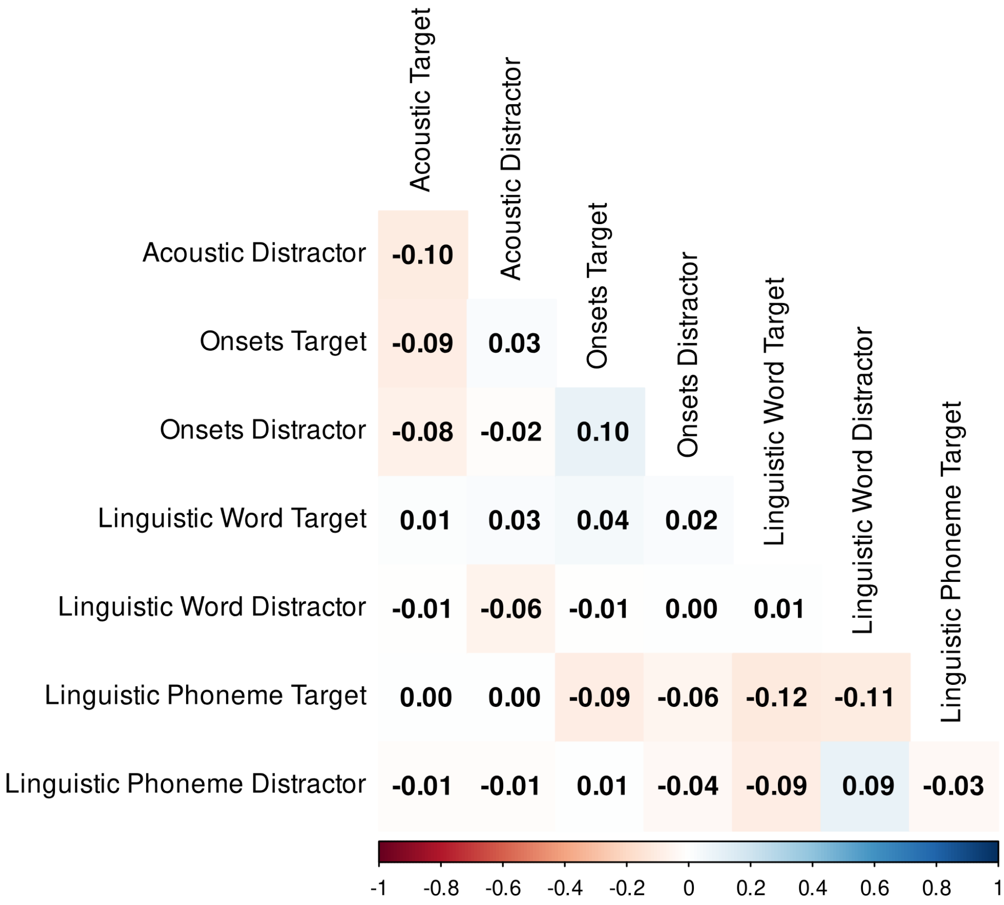


Correlations between the model fits entered into the logistic mixed effects model. All correlations had small magnitudes and we therefore assume no issues related to multicollinearity.

Table S1: Supplementary Model Outputs

| | Estimate | SE | CI | T | p |
|---|---|---|---|---|---|
| Accuracy (incorrect) | -0.14 | 0.07 | -2.06 | [-0.26, -0.01] | .06 |
| Onsets | -0.87 | 0.09 | -10.14 | [-1.04, -0.7] | 0 |
| Phoneme Linguistic | -0.97 | 0.1 | -9.94 | [-1.16, -0.78] | <.001** |
| Word Linguistic | -0.72 | 0.11 | -6.39 | [-0.95, -0.5] | <.001** |
| Stream (Target) | 0.42 | 0.07 | 6.35 | [0.29, 0.55] | <.001** |
| Accuracy*Onsets | 0.15 | 0.17 | 0.85 | [-0.19, 0.48] | .469 |
| Accuracy*Phoneme Linguistic | 0.41 | 0.2 | 2.09 | [0.03, 0.79] | .058 |
| Accuracy*Word Linguistic | 0.27 | 0.23 | 1.2 | [-0.17, 0.72] | .294 |
| Accuracy*Stream | -0.25 | 0.13 | -1.9 | [-0.51, 0.01] | .08 |
| Onsets*Stream | -0.45 | 0.17 | -2.64 | [-0.79, -0.12] | .015* |
| Phoneme Linguistic*Stream | -0.53 | 0.2 | -2.71 | [-0.91, -0.15] | .014* |
| Word Linguistic*Stream | 0.08 | 0.23 | 0.34 | [-0.37, 0.52] | .810 |
| Accuracy*Onsets*Stream | -0.98 | 0.34 | -2.85 | [-1.65, -0.3] | .009** |
| Accuracy*Phoneme Linguistic*Stream | -1.19 | 0.39 | -3.04 | [-1.95, -0.42] | .006** |
| Accuracy*Word Linguistic*Stream | -1.3 | 0.45 | -2.86 | [-2.19, -0.41] | .009** |

Figure S5: Model Outputs for the Distractor Stream

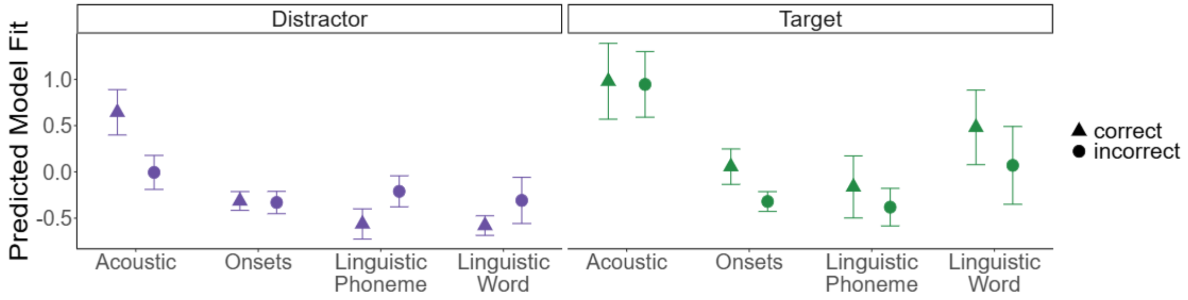


Model fits predicted from accuracy (correct versus incorrect), feature group, stream (target versus distractor), and their interactions.

Table S2: GPT Prompts for target sentence generation

| **Prompts** |
|---|
| Give me 10 sentences with 5-7 words that could stem from a story. |
| Generate 10 everyday sentences with 5-7 words. Only use frequent words and do not use commas. Variate the sentence structure. Do not use question marks or commas. |
| Generate more complex sentences with 5-7 words. Only use frequent words and do not use commas. Variate the sentence structure. Do not use question marks or commas. |

Note. The same prompts were used for the distractor sentences, replacing 5-7 words with 7-10 words.
